## Supplementary Information for "Incompatibility of DFHBI based fluorescent RNA aptamers with particular commercial cell-free expression systems"

**Supplementary Information 1: DNA Sequences (5' – 3')**

T7_F30_Broccoli_Term

```
GTTAGAGAATAGGGATCCGGCGATGCCCGCGAAATTAATACGACTCACTATAGGGTTGCCATGTGTA
TGTGGGAGACGGTCGGGTCCAGATATTCGTATCTGTCGAGTAGAGTGTGGGCTCCCACATACTCTGA
TGATCCTTCGGGATCATTCATGGCAAGCACCGCTGAGCAATAACTAGCATAACCCCTTGGGGCCTCT
AAACGGGTCTTGAGGGGTTTTTTGAATCTGCAGTATGAATTTACTTTAGGCAACTAGAAGGCACAGT
```

FWD_Primer_T7_F30_Broccoli

```
AGAATAGGGATCCGGCGAT
```

REV_Primer_T7_F30_Broccoli

```
GCCTAAAGTAAATTCATACTGCAGATTC
```

T7_F30_iSpinach_Term

```
GTTAGAGAATAGGGATCCGGAATTCGCGGCCGCTTCTAGAGTAATACGACTCACTATAGGGT
TGCCATGTGTATGTGGGAGACGCGACTACGGTGAGGGTCGGGTCCAGTAGCTTCGGCTACTG
TTGAGTAGAGTGTGGGCTCCGTAGTCGCGTCTCCCCACATACTCTGATGATCCTTCGGGATC
ATTCATGGCAAATAACCCCTTGGGGCCTCTAAACGGGTCTTGAGGGGTACTAGTAGCGGCCG
CTGCAGGAATCTGCAGTATGAATTTACTTTAGGCAACTAGAAGGCACAGT
```

FWD_Primer_T7_F30_iSpinach

```
GCCGCTTCTAGAGTAATACGAC
```

REV_Primer_T7_F30_iSpinach

```
GCCTTCTAGTTGCCTAAAGTAAATTC
```

**Supplementary Information 2: 3D Printed Microcuvette adapters**

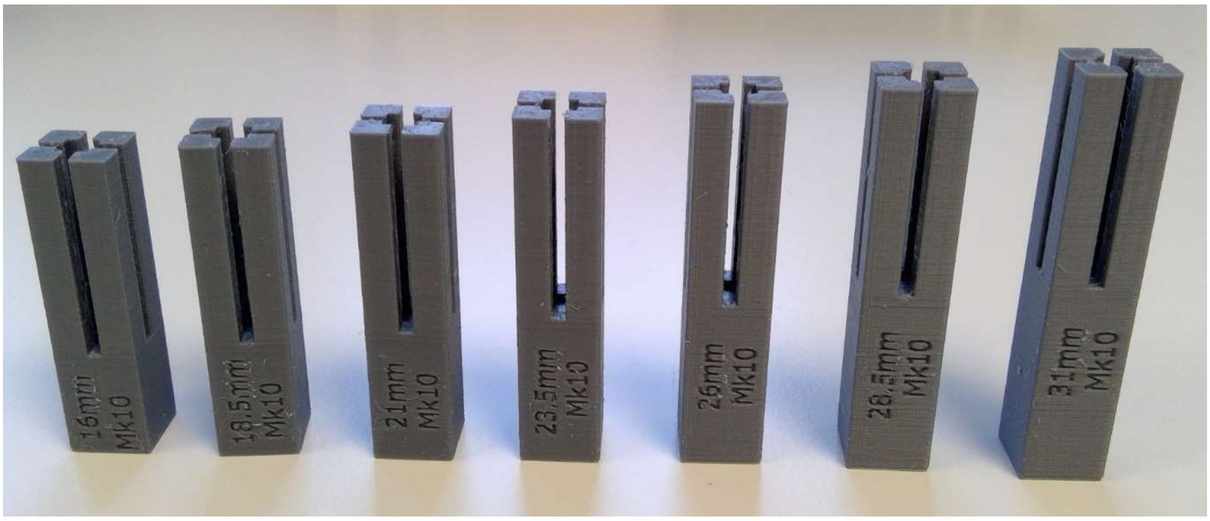


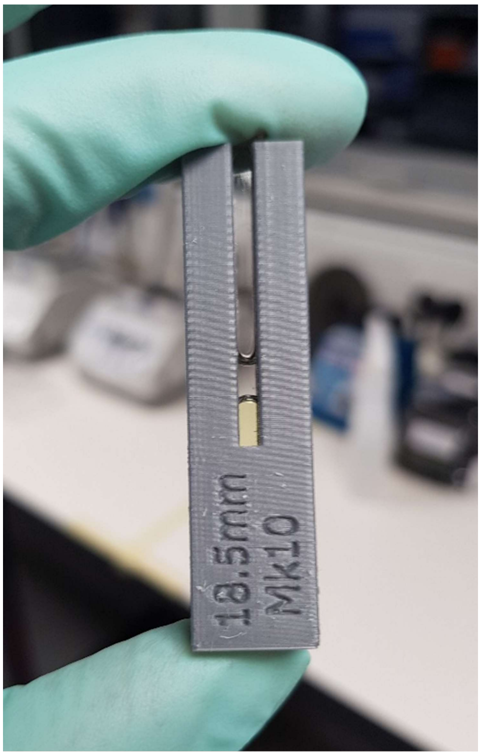


Microcuvette adapters were designed using Autodesk Inventor (v2019.1.1). 18.5mm was found to be the optimal height for the Agilent Cary Eclipse light path. Parts were printed on a Prusa Mk3S using PLA, 0.15mm layer height. Above image shows 100µL reaction mixture + 50µL mineral oil to prevent evaporation.
